# Comparative genomics illuminates adaptive evolution of DVNP with lifestyle and with loss of histone H1 in dinoflagellates

**DOI:** 10.1101/2024.02.09.579734

**Authors:** Jingtian Wang, Hongfei Li, Ling Li, Yujie Wang, Senjie Lin

**Author notes:** Correspondence: Senjie Lin,.

## Abstract

About ten years ago dinoflagellate/viral nucleoprotein (DVNP) was discovered in dinoflagellates, an ecologically important and evolutionarily enigmatic group of aquatic protists. Apparently acquired from a viral origin, the appearance of DVNP coincided with the loss of nucleosome, a rare event in eukaryote evolution. Despite the potential importance of DVNP as the substitute of histones, its evolutionary trajectory and adaptive significance remain elusive. Here, we conducted comparative analyses using existing dinoflagellate genomes and transcriptomes from 26 species ranging from ancestral to later-diverging lineages to investigate the pattern of sequence and structural divergence. Results showed that the functional domestication of DVNP in ancestral dinoflagellates coincided with the loss of histone H1, while subsequent DVNP differentiation was accompanied by the yet another genomic innovation: acquisition of bacterial-originated histone-like protein. Furthermore, our data split DVNP into two major groups: the core DVNP that resembles histone H1 and shows consistently high levels of expression and the non-core DVNP with higher sequence variability and showing lower yet variable levels of expression. In addition, we observed a trend in DVNP evolution tracing that in lifestyle differentiation. This work offers insights into the adaptive evolution of DVNP, laying the foundation for further inquiries of evolutionary drivers and functional innovation of DVNP to enhance our understanding of dinoflagellate evolution and ecological success.

## INTRODUCTION

Dinoflagellates are ecologically important single-celled eukaryotic algae for their vast diversity and major contribution to global marine primary production on the one hand and to harmful algal blooms on the other. Besides, species from the family Symbiodiniaceae in the order Suessiales are indispensable endosymbionts of reef-building corals. Dinoflagellates are cytologically and genomically enigmatic^1–3^. With immense and wide-ranged genome sizes (∼1 to 250 Gbp), dinoflagellate chromosomes are permanently condensed^4–6^. Recent third-generation genome sequencing coupled with HiC linkage analysis revealed a 2-D structure of chromosomes in Symbiodiniaceae^7,8^. In this model, chromosomes are composed of highly interactive DNA blocks, named topologically associated domains (TADs), which harbor discrete cis- and trans- oriented unidirectional gene clusters. Within each TAD lies the switch point of the cis- and trans- gene clusters (diverging strand switch region or dSSR), which were postulated to harbor promoters for gene transcription, whereas TAD boundaries (convergent SSR or cSSR) feature low gene density and harbor transcriptionally less active genes, which encode long-lived RNA and stable proteins needed for chromosome structure maintenance^1^. Besides, the 2-D model revealed a high gene density at chromosome ends, which probably allows access to genes by transcription machinery for gene expression while the chromosomes remain generally condensed^1^.

Another peculiar feature of dinoflagellate nuclei is the lack of nucleosome^4–6^. Nucleosomes are the fundamental units of chromatin in eukaryotic organisms, consisting of a complex of core histone proteins (H2A, H2B, H3, and H4) surrounded by DNA helix and connected by histone H1. Histones not only facilitate DNA condensation but also, with post-translational modifications, play a crucial role in genome regulation^9–11^. Thus, histone proteins are highly conserved and indispensable for ensuring the growth and development of eukaryotic organisms^12^. It was long believed that dinoflagellates had lost histone genes^13^.

The notion changed when omics surveys revealed the presence and transcription of core histone genes in dinoflagellates^14^. Despite this, the encoded histone proteins have never been detected, likely due to very low abundances, suggesting that these histones probably do not play a major role in packaging dinoflagellate genomes^14–16^. Gornik et al. (2012) discovered Dinoflagellate/Viral NucleoProtein (DVNP) in the parasitic (ancestral) dinoflagellate *Hematodinium*, and found that the loss of nucleosomal DNA condensation in dinoflagellates coincides with the appearance of DVNP. DVNP is highly positively charged, is nucleus-localized, binds to DNA with similar affinity to histone proteins, has several phosphorylation states, and is found only in dinoflagellates besides marine large DNA viruses^4^. These nucleoproteins are hence hypothesized to have been transferred from viruses to ancestral dinoflagellate with typical chromatin, eventually functionally replacing histones as the chromatin packaging proteins^17,18^. In support of this hypothesis, when introduced into *Saccharomyces cerevisiae*, DVNP expressed in the budding yeast was found to remodel chromatin and preferentially associated with nucleosomal regions^12^.

Despite its potential importance in dinoflagellate nuclear function, DVNP is in general poorly understood and understudied. A systematic genetic study of DVNP is needed to illuminate its evolutionary and functional differentiation among lineages and decipher the role of DVNPs in adaptive evolution of dinoflagellates. In this study, we investigated the gene copy dynamics and sequence divergence of DVNP across an evolutionary spectrum of dinoflagellates by using newly generated and existing omics data for 26 species that represent the major lineages of dinoflagellates. We also analyzed the structure, evolutionary characteristics, and expression patterns of DVNP in these lineages.

Furthermore, we compared DVNP to histones and histone-like proteins (HLPs). Results revealed lifestyle-dependent DVNP copy number variation, strong purification evolution, apparent functional innovation (in some DVNPs), and evidence of evolutionary trajectory from initial domestication to differentiation selected for due to bacterial-derived HLP transfer.

## RESULT

### Lifestyle-dependent trend of duplication and divergence of DVNP

We generated a series of transcriptomes from *Prorocentrum shikokuense* and collected data on other species from sequencing projects (Table S1). In total, we identified 719 DVNP genes (Supplementary Data set 1) from 26 dinoflagellates that were grouped into four ecotypes: parasitic (2 species), free-living and bloom-causative (9 species), free-living and non-bloom causative (3 species), and symbiotic (12 species). We also obtained 36 viral homologs of DVNP (VNP) from Uniprot. For *Hematodinium* sp. and *Breviolum minutum*, we identified 13 and 18 DVNP genes, respectively, consistent with previous studies^16,19^, indicating that our identification pipeline is reliable.

We mapped the genome information and the number of identified DVNP genes to a genome-based phylogenetic tree (Fig 1A). The topology of the tree is consistent with previously published phylogenies, confirming its reliability^18^. *Perkinsus marinus* was included in the tree as the outgroup because it is the closest alveolate relative to dinoflagellates and has no DVNP but has classical histones-based nucleosomes. All 26 dinoflagellates have DVNP genes but their copy numbers vary with species. Although larger genomes usually mirror more gene duplications^3,20^, the number of DVNP genes does not scale with genome size (Fig 1A). Between the two major parasitic genera from the order Syndiniales, *Hematodinium* has a 40-fold larger genome and ∼3-fold greater gene repertoire relative to *Amoebophrya*. However, the two species have similar DVNP gene copies (10 and 13, respectively). To examine whether DVNP copy number has a consistent relationship with ecotype or lifestyle, we compared symbiotic with free-living bloom-causing species. Results show that the numbers of DVNP genes are more similar among the symbionts (11 to 21) than among bloom-causative free-living species (10 to 59). The Arctic (CCMP2088) and Antarctic (CCMP1383) strains of *Polarella gracialis* both have the highest and similar copy numbers (129 and 161 for the Arctic and Antarctic strains, respectively), indicating a similarly remarkable DVNP gene family expansion in them despite their extreme geographic isolation (Fig 1A).

**Fig 1.**
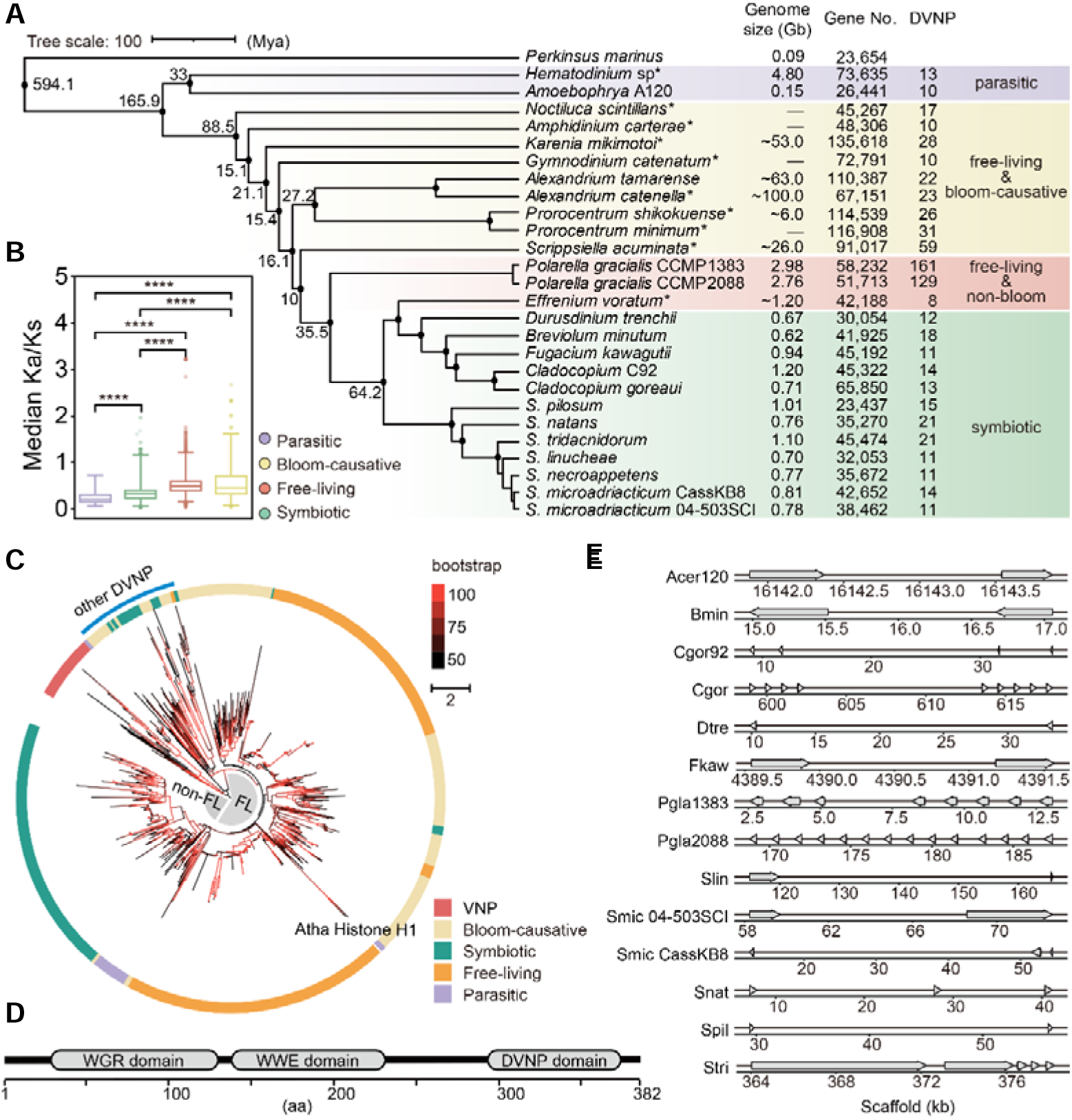
Phylogenetic trend of DVNP gene copy number and structural features in 26 dinoflagellates. **(A)** Evolutionary dynamics of genome size, coding gene number, and DVNP gene copy number mapped to multiprotein phylogeny of dinoflagellates with *Perkinsus marinus* as the out group. Nodes in the tree show lineage divergence time based on TIMETREE database (Mya, million years ago). Asterisks indicate data based on transcriptomes. **(B)** Boxplot of Ka/Ks ratio from all identified DVNP domain-containing genes in 26 species. Asterisks indicate strong significance (Mann–Whitney *P*L≤L0.0001). Boxplots show the Q1 and Q3 (the 25th and 75th percentile or the IQR), with the median in the center and the dots denoting outliers (<5% or >95%). **(C)** Maximum-likelihood DVNP phylogeny (IQ-Tree) revealing evolution of the DVNP and affiliation of *A. thaliana* histone H1 within DVNP. Ultrafast bootstraps at branches are shown in the color bar (from 50 in black to 100 in red). The topology tests of Atha histone H1 in the DVNP tree rejected several hypotheses tests, all tests performed 10000 resamplings (Table S2). FL: free-living species (bloom-causing or not); non-FL: non-free-living species (symbiotic and parasitic species). **(D)** Schematic structure of DVNP with two additional domains (WWE and WGR) in the ‘other DVNP’ clade. **(E)** Scaffold coordinate and arrangement of DVNP genes in dinoflagellates genomes. Arrows depict gene orientation. Species name abbreviation: Acer120, *Amoebophrya ceratii* 120; Bmin, *Breviolum minutum*; Cgor92, *Cladocopium* C92; Cgor, *Cladocopium goreaui*; Dtre, *Durusdinium trenchii*; Fkaw, *Fugacium kawagutii*; Pgla1383, *Polarella gracialis* CCMP1383; Pgla2088, *Polarella gracialis* CCMP2088; Slin, *Symbiodinium linucheae*; Smic_04-503SCI_03, *Symbiodinium microadriacticum* 04-503SCI.03; SmicCassKB8, *Symbiodinium microadriacticum* CassKB8; Snec, *Symbiodinium necroappetens*; Snat, *Symbiodinium natans*; Spil, *Symbiodinium pilosum*; Stri, *Symbiodinium tridacnidorum*.

DVNP sequence evolution also appears to be associated with lifestyle. Free-living (bloom-causing or not) species have higher ratios of non-synonymous to synonymous substitutions (Ka/Ks) in the DVNP genes than species of non-free living (parasitic or symbiotic) lifestyle, while no significant difference between species within the free-living group. As higher Ka/Ks signals sequence divergence more likely to be selected for than neutral variations, our result suggests that the DVNP sequence divergence was under selection pressure either resulting from, or co-acting on, lifestyle (Fig 1B). Furthermore, we found that the clustering of DVNPs is related to lifestyle differentiation (free-living and non-free-living) (Fig 1C). The only exception is that there is one clade (named ‘other DVNP’) that contains free-living and symbiotic species, in which the DVNP contains two additional domains (WWE and WGR) in all symbiotic and some bloom-causing free-living species within this clade. The WWE and WGR domains are highly conserved across the species that contain them (Fig 1D and Fig S1).

Based on published dinoflagellate genome assemblies, we noted that DVNP genes are unidirectionally encoded and tandemly arranged (Fig 1E and Fig S2), as the case of many dinoflagellate genes^21,22^. The two strains of *P. gracialis* have the greatest number of tandem repeats of DVNP (up to 14 copies found in one scaffold, Fig 1E and Fig S2). In contrast, tandem repeats of DVNP seem to be rare in parasitic species, as we only found two scaffolds in *Amoebophrya ceratii* with tandem repeats of DVNP, and each scaffold has no more than three DVNP copies (Fig 1E and Fig S2).

### Differentiation of DVNP into two subgroups

Dinoflagellate nucleus harbors the vertically inherited ancestral core histones, the virus-originated DVNP, and bacterium-originated histone-like proteins. This multi-origin of nuclear proteins raises a question of how other nuclear proteins co-evolved with DVNP, To explore this question with existing data, we identified histones and HLPs from the 26 dinoflagellates covered in this study (Supplementary Data set 2), and used them along with DVNP to construct a similarity-based clustering (Fig 2A). The sequence distance-based cladogram shows a clear separation of histones, HLPs, and DVNPs, and the ubiquity of core histones (H2A/B, H3, H4) in dinoflagellates, consistent with previous reports on various species of dinoflagellates^14,15,23^. The clustering pattern showed that DVNPs branch from viral homologs of DVNP (VNP) and form multiple subclades. The large sequence divergence of DVNPs between species indicates a rapid evolution of DVNP (Fig 2B).

**Fig 2.**
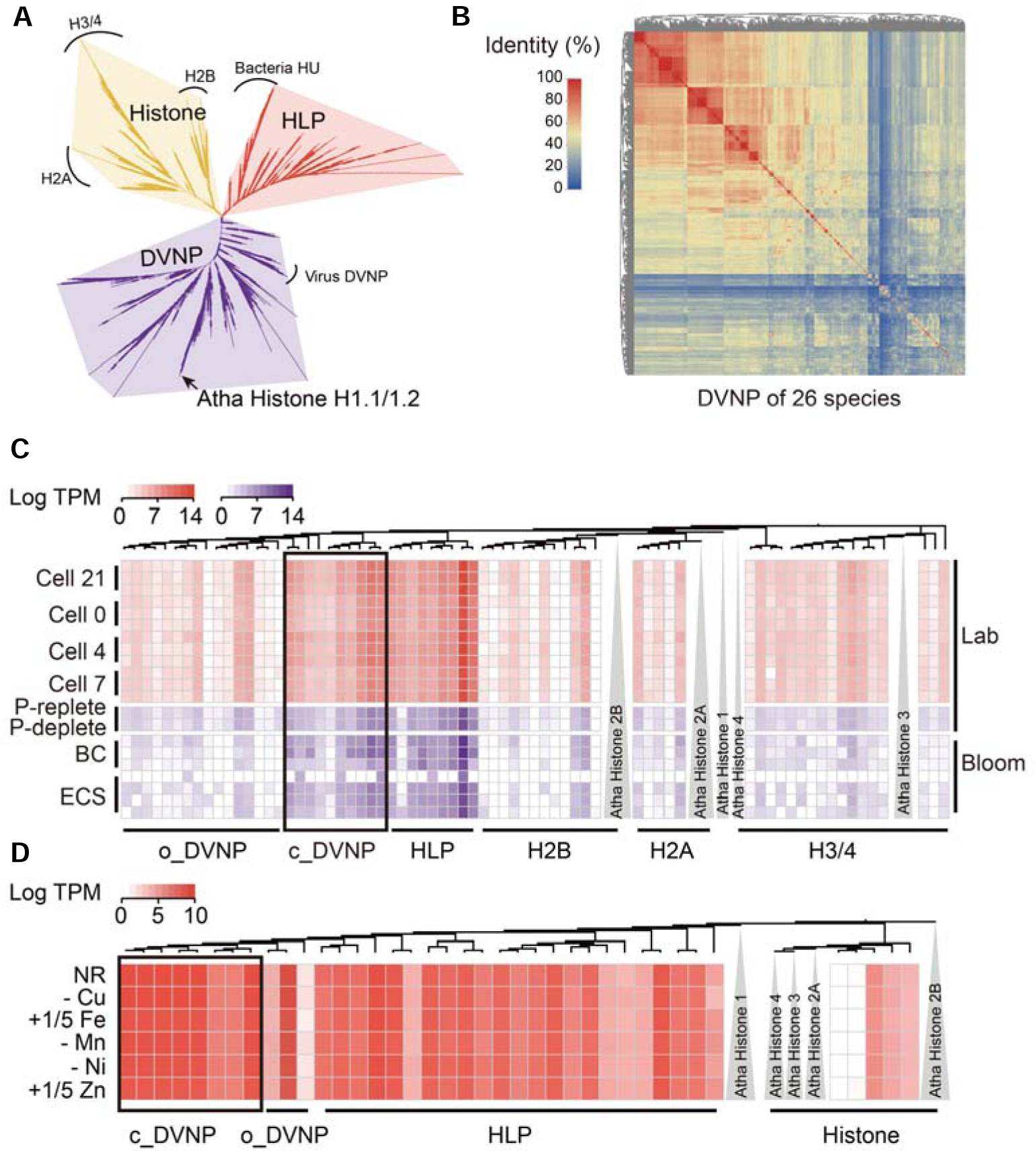
The evolutionary divergence of DVNP in sequence and expression pattern. **(A)** Maximum likelihood phylogeny (IQ-Tree) of DVNPs, histones and HLPs. All types of histones in *A. thaliana* are clustered with dinoflagellate histones, except *A. thaliana* histone H1, which is clustered with DVNP (arrow). Arches mark clusters. **(B)** Pairwise identity hierarchical clustering pattern of all dinoflagellate DVNPs. **(C-D)** Phylogeny of DVNP, histones and HLP of *P. shikokuense* **(C)** and *F. kawagutii* **(D)** and their expression profiles across different conditions. Cell cycle (21h, 0h, 4h, 7h) and phosphorus treatment (P-replete, P-replete) in the lab, two *P. shikokuense* blooms in China Sea (BC, Baicheng, Xiamen; ECS, East China Sea, Zhejiang). Data of *F. kawagutii* were derived from our SAGER Database (http://samgr.org.cn/). Abbreviations: o_DVNP, other (non-core) DVNP; c_DVNP, core DVNP; NR, nitrate replete; -Cu, without Cu; +1/5 Fe, 50 nM, 1/5 of its normal concentration; -Mn, without Mn; -Ni, without Ni; +1/5 Zn, 2 nM, 1/5 of its normal concentration.

The evolutionary divergence of DVNP does not only occur in sequence, but also in expression pattern. Based on transcriptomes of various dinoflagellates under different conditions, the expression profile divides DVNPs into two groups, even within species (Fig 2C, D and Fig S3). One of them was highly expressed regardless of growth conditions of the cultures, herein named ‘core DVNP’. The other group is composed of all remaining subclades of DVNP and was expressed at low or undetectable levels under any growth condition (Fig 2C, D and Fig S3), which corresponds to the ‘other DVNP’ cluster in the cladogram (Fig 1C). The high expression of core DVNP seems to be a shared trait with histone-like proteins (HLP), which were consistently found to be highly expressed in various species under different conditions both in previous studies^3^ and the present study (Fig 2C, D). Interestingly, HLP is believed to be a bacteria-originated substitute of histones in dinoflagellates^24^.

### Similarity of core DVNP to *Arabidopsis thaliana* histone H1

Because core histones interact with histone H1 in nucleosome formation, we examined if this gene is present in the dinoflagellate omic dataset, and if so whether it diverged from typical histone H1 along with the loss of nucleosome. Strikingly, our exhaustive search into all the existing data from the 26 dinoflagellate species, encompassing ancestral and advanced taxa and data from public databases (NCBI, UniProt) as well as our in-house Symbiodiniaceae and Algal Genome Resource database (SAGER), yielded no histone H1 (with H1 domain PF00538) in any of the species. In addition, we have carefully reviewed the main reports on dinoflagellate histone H1 and cannot confirm its accuracy^23,25–27^. This indicates that histone H1 was probably lost early in dinoflagellate evolution. We could not rule out the possibility that dinoflagellates contain histone H1 genes but their sequences have diverged too far to be recognizable, we think the likelihood is small. Interestingly, when using DVNP sequence in the same search, it not only hit all the dinoflagellate homologs but also, albeit with weak E value, plant histone H1. To compare DVNP and histone H1 for sequence similarity, we included both proteins in the clustering analysis. From this inclusive cladogram (Fig 1C and Fig. 2A), histone H1 of the plant model *A. thaliana* was clustered with the ‘core’ subgroup of DVNPs. The clade includes core DVNPs from *P. shikokuense*, *F. kawagutii* and the other 21 species as well as *A. thaliana* histone H1 (Fig 1C and Fig 2A).

Furthermore, core DVNPs show a significantly higher sequence similarity to *A. thaliana* histone H1 than to non-core DVNPs or VNPs (Fig. 3A). Specifically, *A. thaliana* histone H1 shares ∼10% identity (i.e. 9.2% in *P. shikokuense*, 10.3% in *F. kawagutii*) with the core DVNPs (Fig 3B, C) but only 2.9% or less with non-core DVNPs in the cases of *P. shikokuense* and *F. kawagutii* (Fig S4). An alignment analysis showed highly conserved amino acid residues between core DVNPs and the histone H1 domain, particularly at the potential methylation residue lysine, but less conserved sites between core DVNP and VNPs (Fig 3B, C and Fig S5, S6). In addition, the core DVNPs are more conserved to each other at some residue sites than to the canonical DVNP domain (pfam domain PF19060) (Fig. 3D).

**Figure 3.**
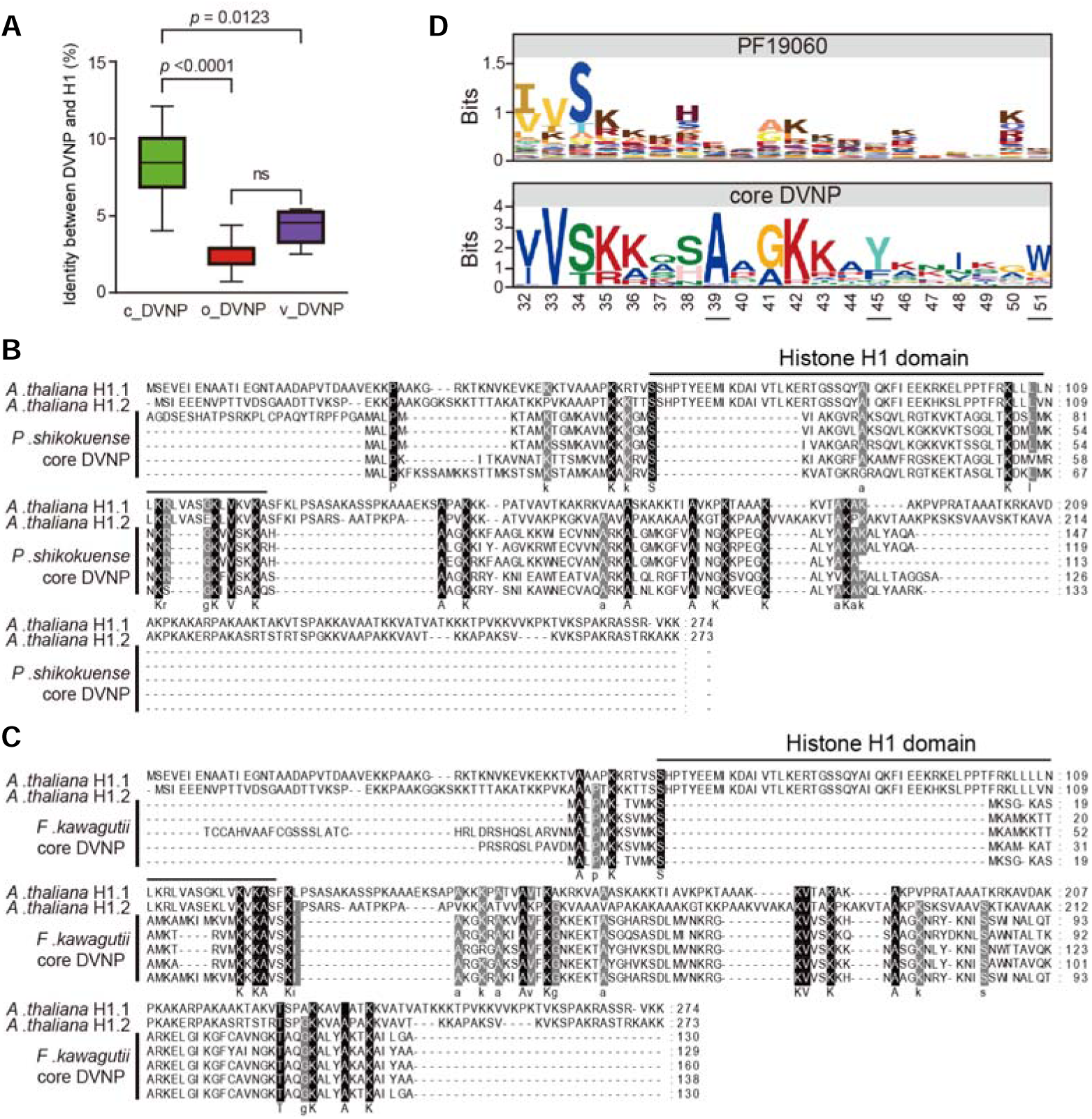
Identity and conservative domain of core DNVP in dinoflagellates. **(A)** Box plot of the identity among DVNPs and *A. thaliana* histone H1. Number indicates Mann–Whitney *P*-value; c_ DVNP, core DVNP; o_ DVNP, other DVNP; VNP, viral homolog of DVNP; ns, not significant. **(B-C)** Multiple alignments of *A. thaliana* histone H1 and core DVNP genes of *P. shikokuense* **(B)** and *F. kawagutii* **(C)**. Small letters in the consensus line indicate conserved residue (conserved percent > 80%). *A. thaliana* H1.1, H1.2 (Uniprot, P26568, P26569). **(D)** Graphical motif representation showing conserved amino acid residues of canonical DVNP domain and core DVNP.

The coincidence of core DVNP appearance with histone H1 disappearance is interesting and unsuspected. Along with the substantial similarity between core DVNPs and plant H1, the finding suggests that DVNP has diverged radically since its acquisition from its viral source, with the core subgroup evolving towards resemblance to histone H1 during its ‘domestication’ in the dinoflagellate host.

## DISCUSSION

Here, we utilized extensive dinoflagellate omics data to systematically identify and characterize DVNP that are unique in dinoflagellates and aimed to shed light on the evolutionary trajectory and potential functions of DVNP in dinoflagellate. We found evidence that DVNP copy number variations (CNV) and sequence divergence was coincided with dinoflagellate lifestyle differentiation. Furthermore, we found that DVNPs evolved fast, differentiating into two distinct groups that are expressed at different levels. In addition, one of the DVNP subgroups (core DVNP), which consistently displays high expression levels in all dinoflagellate species grown under various conditions, resembles plant histone H1. These findings still provide valuable insights into the complex nuclear evolution in dinoflagellates.

### Divergence of DVNP among dinoflagellates with lifestyle

DVNP copy number appears to be relatively stable in non-free-living species (parasitic and symbiotic) but shows a trend of contraction compared to free-living species, indicating a closer relationship of DVNP copy number to lifestyles than genome size.. Compared with symbiotic species, DVNPs in *P. glacialis* exhibit high copy number and frequent tandem repeats. Gene tandem repeats are common in dinoflagellates^22,28^. Considering the complex external environment, the expansion of the DVNP gene family may be a mechanism to quickly increase or regulate their expression to enable swift responses to environmental fluctuation, and this expansion and tandem repeats drive the evolution of DVNP^21^.

It is interesting that the evolution rate of DVNP also varies with lifestyle. Our analysis results showed a sign of purification selection (Ka/Ks<1) for the majority of DVNP, an evolutionary characteristic shared by eukaryotic histones. The purification evolution suggests that the function of DVNP is crucial throughout the dinoflagellate phylum^29–31^. However, DVNP exhibits differential evolutionary rates between dinoflagellates of different lifestyles, except within the group of free-living dinoflagellates (between non-bloom and bloom-causative species). A small subset of DVNP shows positive selection for divergence (Ka/Ks > 1), some of them acquiring two additional domains (WWE and WGR), indicating functional adaptive evolution in this group of DVNP. This sequence divergence and potentially functional differentiation of this subset is confirmed through sequence identity and gene expression patterns, which will be discussed below.

### Rapid DVNP evolution and differentiation of a histone H1-like subclade

The 13 DVNPs previously reported in *Hematodinium* sp. exhibit high sequence identity^19^, but our data indicate that not all species share the same conservativeness. Some studies have revealed diversity in DVNP sequences^8,16^, yet not until now has a systematic examination for different lineages of dinoflagellates been conducted. Based on comparative analyses of genomes and transcriptomes, we found evidence of rapid DVNP gene sequence divergence. The evolution led to its differentiation into two groups: a monophyletic subclade (core) in the distance-based cladogram that contains plant histone H1 and a multi-subclade group (non-core). Furthermore, our data show that in the evolutionary history of dinoflagellates, while the core-DVNP was ‘ancestral’, the non-core DVNP first appeared in the Amphidiniales lineage and later widely spread to later-diverging lineages (Fig. 4). Compared to non-core DVNP, core DVNP is more similar to (∼10% similarity) eukaryotic histone H1.

**Fig 4.**
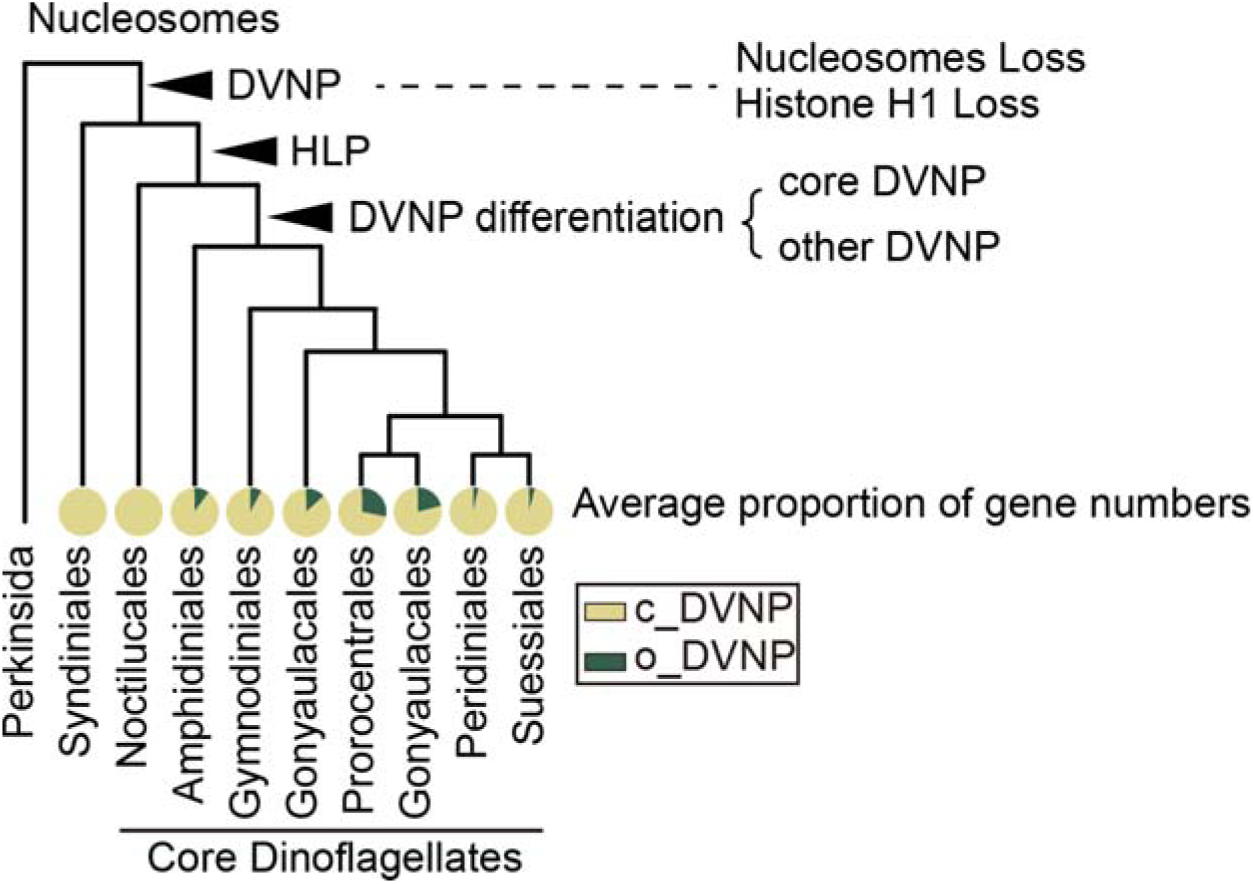
Model for character evolution in dinoflagellates. Model depicting a consensus phylogeny of dinoflagellates and the histone-related gene traits (arrowheads). The pie chart is based on data from 26 species in this study. c_DVNP, core DVNP; o_DVNP, other (non-core) DVNP.

The finding of the histone H1-like DVNP is reminiscent of the documentation of histone-like protein (HLP). Based on a study of HLP (also known as HCc) in *Crypthecodinium cohnii*, which also possesses DVNP differentiation (Fig S7), dinoflagellate HLPs appear to have originated from a DNA-binding protein of bacteria (HU), and they share ∼10% sequence identify with histone H1 of eukaryotes^24^. HCCs have been postulated to represent a continuum in the evolution from the prokaryotic HU to the eukaryotic histone H1^24^. Based on our data of sequence similarity, core DVNP may also be an intermediate in the evolution continuum from VNP (viral source of DVNP) to eukaryotic histone H1. The considerable sequence similarity of core DVNP to eukaryotic histone H1 observed in this study suggests that DVNP has undergone an adaptive evolution, which might have led to neofunctionalization, possibly driven by the absence of histone H1, and as part of the ‘domestication’ in the dinoflagellate intracellular environment since the acquisition of the viral gene.

### New evolutionary complexity in the nuclear evolution of dinoflagellate

A new interesting twist has emerged to the currently known trajectory of dinoflagellate nucleus evolution (Fig 4). The acquisition of DVNP by ancestral dinoflagellates (parasitic taxa) has been associated with the loss of nucleosome^19^. Now with the absence of histone H1 in ancestral and advanced lineages of dinoflagellates revealed in our study, the loss of histone H1 was another part of the evolutionary event. Although proteins of canonical histones have never been detected^15^, their encoded genes and transcripts occur throughout the tree of dinoflagellates^14,15^. Therefore, it can be inferred that the loss of nucleosome is probably more directly associated with the loss of histone H1 and gain of DVNP than the functional ‘fall’ of histone genes, the latter of which might rather be a derived trait. Further research is required to more rigorously examine this scenario.

The discovery of HLP in dinoflagellates^32–34^ has filled the void of dinoflagellate nuclear architecture left by the functional fall of histones, particularly as HLP appears to be capable of condensing DNA into liquid crystaline state^35^ that characterizes dinoflagellate chromosomes. A study also showed that HLPs have a higher affinity for binding to single-stranded DNA compared to double-stranded DNA^36^, and immunofluorescence in *C. cohnii* showed its localization at the periphery of chromosomes, where active transcription occurs, suggesting a role in gene expression regulation^36^. Our analysis, consistent with previous reports^18^, indicates that HLP was acquired roughly when *Noctiluca* emerged (Fig. 1A, Fig. 4). This major horizontal gene transfer event (from a bacterial source) in dinoflagellates occurred after DVNP acquisition (from a viral source), but was followed by, and possibly drove, the subsequent rapid divergence of DVNP, as suggested by our data.

DVNP has been diversified radically (Fig. 2). Heterologous expression of one DVNP sequence in yeast showed that it binds to DNA and has functional conflict with core histones^12^. Data reported in the present study and previously in concert suggest that core DVNP may possibly functionally complement the lost histone H1 and play a role in organizing chromosomes in dinoflagellates. The non-core DVNP, however, might be involved in regulating transcriptional activities, the role also implicated for HLP. This progressive evolutionary trajectory of dinoflagellate nuclear proteome, involving acquisition and divergence of DVNP and HLP and underlying regulatory circuitry of their functions remains to be experimentally examined and completely deciphered.

## CONCLUSION

The dinokaryon stands out among eukaryotic nuclei in complexity and enigma. The occurrence of viral-originated DVNP coincided with significant changes in nuclear characteristics, including genome enlargement and chromosome condensation as well as loss of histone H1 and nucleosome, throughout the dinoflagellate evolutionary history. Unveiling the role of DVNPs in adaptive evolution is crucial for comprehending the peculiar nuclear features of dinoflagellates. This study shed light on the previously not fully recognized complexity in dinoflagellate nuclear evolution involving serial events involving loss of nucleosome, histone H1, and acquisition of HLP, as well as diversification of DVNP. The evolution of DVNPs after acquisition by ancestral dinoflagellates may be linked to the loss of histone H1, and subsequent DVNP differentiation might have been influenced by bacterial-derived HLP acquisition. This work has provided the first broad insight into the genomic properties of DVNP, and the findings will be helpful for in-depth interrogation of the evolution and functional innovation of DVNP along with interacting nuclear proteins to better understand how the unique dinokaryon confers dinoflagellates their competitiveness and ecological success.

## MATERIALS AND METHODS

### Construction of the complete transcriptome dataset of *Prorocentrum shikokuense*

*P. shikokuense* was obtained from the Culture Collection Center of Marine Algae, Xiamen University, and was cultured in L1 medium. To achieve the most complete transcriptome of *P. shikokuense* as practically feasible, cultures were grown under four conditions (light changes, nutrient changes, growth stages, and temperature changes; see Table S3 for detailed information) each in triplicate. This gave a total of 25 cultures. About 10^8^ cells were collected from each culture daily by centrifugation at 5,000 rpm, 4 °C, for 10 min. The cell pellets were immediately suspended in 1 mL TRIzol reagent (Invitrogen, Carlsbad, CA, USA), vortexed thoroughly, and stored at –80°C for subsequent RNA extraction. RNA extraction was carried out with a Direct-zol RNA MiniPrep kit (Zymo Research, USA) and quantified using a NanoDrop ND-2000 spectrophotometer (Thermo Scientific, Wilmington, DE, USA). All RNA samples were pooled for sequencing using PacBio ISO-Seq and BGISEQ-500 Denovo polyA mRNA workflow (BGI Genomics Co., Ltd.). The sequencing depth was set to achieve 14.7 Gb and 20.3 Gb data output, respectively, totaling 35.0 Gbp. These included triplicate RNA sampling for four diel time points (24:00, 4:00, 7:00, 9:00), and sequenced following the same procedure as above, with 1 Gb data generated for each of the 12 samples. After clustering and error correction using Quiver (Quality Values ≥ 0.98)^37^, and de-redundancy using CD-HIT (-c 0.98 -T 6 -G 0 -aL 0.90 -AL 100 -aS 0.98 -AS 30)^38^, 233,334 unigenes were generated, with N50 of 1,548 bp.

### Gene identification in dinoflagellates

In addition to the comprehensive transcriptome of *P. shikokuense* generated in this study, other dinoflagellate data came from published studies, including genomes and transcriptomes (Table S1). For genome data, we designed a pipeline to identify DVNP genes with efforts to reduce potential chimeric genes. In brief, we collected DVNP sequences from Uniprot^39^ and NCBI^40^, and after checking domain integrity, we kept high-quality DVNP sequences as a reference. Then, Blastall v2.2.26^41^ was used to align genome data to the reference. After extending 2000bp upstream and downstream of the aligned genome segments, GeneWise2^42^ was used to *de novo* predict the gene coding region. Predicted sequences were checked for integrity of the DVNP domain through hmmsearch v3.1b2 (c-Evalue and i-Evalue both < 1e-5)^43^ and CDD (Evalue < 1e-5)^44^, and double-checked through the SAGER database by Diamond v2.0.9.147 ^45^. Eligible sequences were assigned as the high-confidence DVNP genes for each species. Detailed parameters can be viewed through scripts at: https://github.com/WJT0925/DVNP-gene-identify.

For transcriptomes, we first searched the above-mentioned reference using Blastp (1e-5)^41^. The aligned sequences were searched for the presence of the DVNP domain using hmmsearch (c-Evalue and i-Evalue both < 1e-5) and CDD (Evalue < 1e-5). The sequences resulting from the search were subjected to de-redundancy using CD-HIT (-c 0.95 -aS 0.9 -d 0 -M 0) to generate a unigene dataset (Supplementary Data Set 1).

Histone genes of the plant model *Arabidopsis thaliana* were obtained from SwissProt, and bacterial HU-like genes were from a previous study^18^. Histone and histone-like protein genes from 26 dinoflagellates were identified in the same method as DVNP identification (Supplementary Data Set 2).

### Ka/Ks and phylogenetic analysis

Paired-align identity and Ka/Ks substitutions between gene pairs were calculated using TBtools^46^. OrthoFinder v2.5.4^47^ was used to infer the species phylogeny among the 26 dinoflagellates and the early-branching *Perkinsus marinus*.To construct the species tree, orthologous genes were obtained from all-against-all blastp (Evalue 1e-5), clustered, and used for gene tree inference. Sequences were aligned using Muscle^48^. To construct DVNP tree and the comprehensive phylogenetic tree of DVNP along with the histones and histone-like proteins, outgroups were collected: viral homologs of DVNP from Uniprot (VNP); histone protein of *A. thaliana* from Uniprot reviewed data; bacterial HU-like protein from reference^18^. All Maximum Likelihood phylogenetic trees in this study were constructed using IQ-TREE2 (1,000 ultrafast bootstraps)^49^. The topology test of Atha histone H1 in the DVNP tree were hypotheses two conditions, histone H1 clustered in other DVNP or histone H1 out of DVNP cluster, then test using IQ-TREE2 (-n 0 -zb 10000 -au).

### Expression analysis

Multiple transcriptomic datasets were used in this analysis. Metatranscriptomes of two *P. shikokuense* blooms were used^50^. One of the blooms occurred in 2014 at Yangtze River Estuary, East China Sea (ECS; data covering pre-bloom sample of T0 and bloom period samples of T1, T2, T3)^51^. The other occurred in the same month and year at Baicheng Beach, Xiamen Harbor (BC; data covering samples of 11:00 pm on May 6 and 5:00 am and 1:00 pm on May 7)^52^. The expression (TPM) of *F. kawagutii* across different conditions was obtained from Symbiodiniceae and Algal Genomic Resource (SAGER, http://sampgr.org.cn/index.php)^53^. Samples of *S. acuminate* representing different cell stages were obtained from GenBank’s Sequence Read Archive (SRA) database, PRJNA744760^54^. Expression (TPM, Transcript per million) of all transcriptomes was calculated by RSEM (version v1.3.3, parameter: default)^55^.

### Core DVNP group identification and motif prediction

To determine the phylogenetic relationship of these genes along with the histones and histone-like proteins in all the species, a comprehensive phylogenetic tree was constructed. The clade composed of *A. thaliana* histone H1 was identified for each species and designated as core DVNPs based on properties described in the results (Supplementary Data set 1). Conserved domains in core DVNPs were identified using MEME SUITE (-mod zoops -nmotifs 1 -minw 6 -maxw 50 -objfun classic -markov_order 0)^56^.

## AVAILABILITY OF SUPPORTING DATA

All the raw data of *P. shikokuense* culture-based transcriptomes have been deposited at the NCBI Sequence Read Archive (SRA) database (http://www.ncbi.nlm.nih.gov/) under BioProject no. PRJNA990325. The processed transcriptomic data of *P. shikokuense* is publicly available on the Symbiodiniceae and Algal Genomic Resource website (SAGER, http://sampgr.org.cn/index.php). Other data used in the study have been published and publicly available at NCBI (see Table S1).

## AUTHOR CONTRIBUTIONS

Senjie Lin and Jingtian Wang conceived the work. Jingtian Wang and Hongfei Li carried out the experiments and data analysis. Ling Li and Yujie Wang provided technological and logistic support. Jingtian Wang wrote the manuscript. Senjie Lin reviewed the manuscript. All authors participated in revising the manuscript and agreed to the final submitted version.

## ACKNOWLEDGEMENTS

This work was financially supported by the National Key Research and Development Program of China (Grant No. 2022YFC3105301), National Natural Science Foundation of China (Grant No. 42276096). Lin was partially supported by the Gordon and Betty Moore Foundation (Grant No. #4980.01).

## Supplement Fig & Table

**Fig S1.**
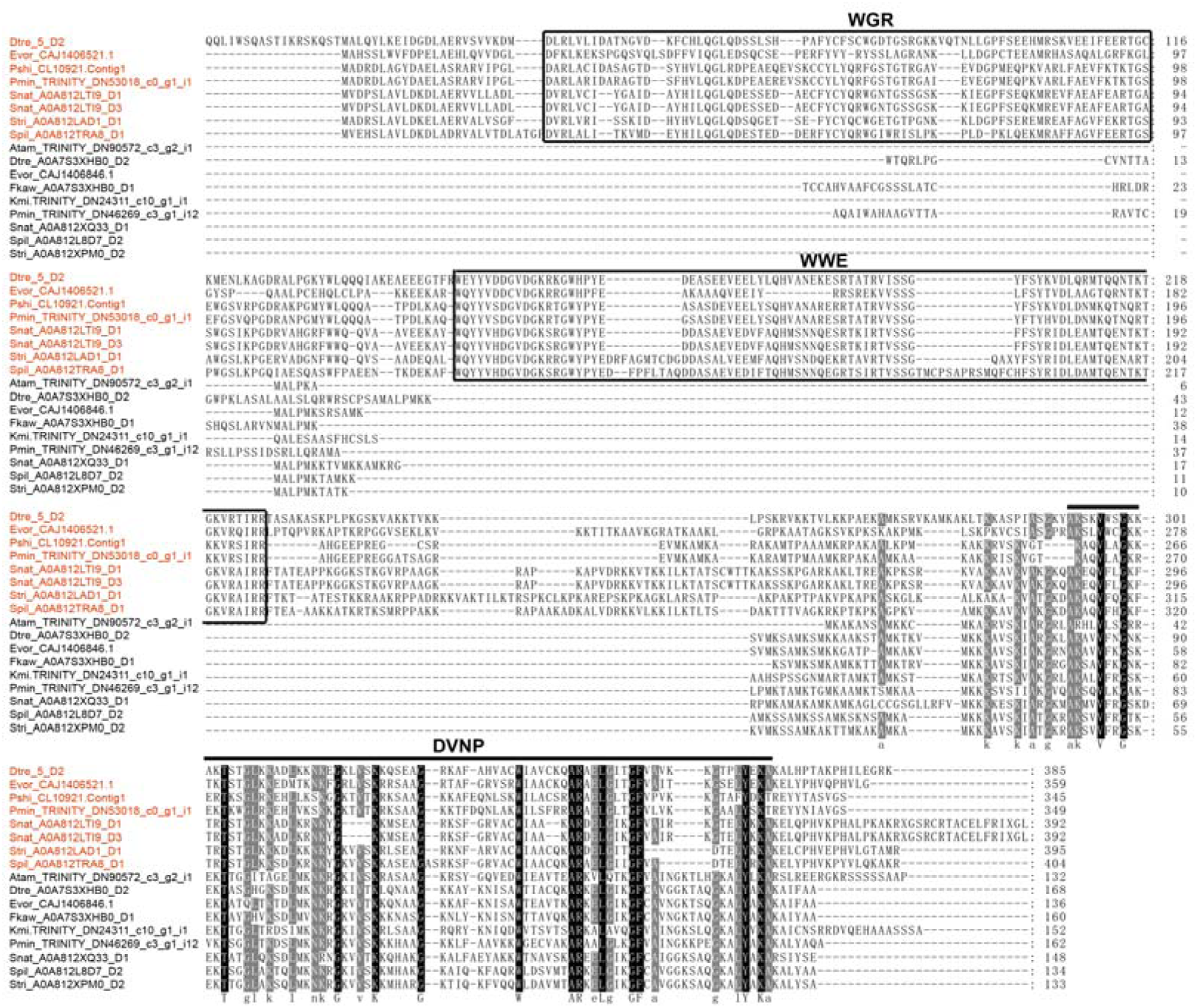
Alignment of representative sequences between DVNPs containing the WGR and WEE domains and those lacking these domains. The red label indicates DVNPs containing WWE and WGR domains. Small letters in the consensus line indicate conserved residue (conserved percent > 80%).

**Fig S2.**
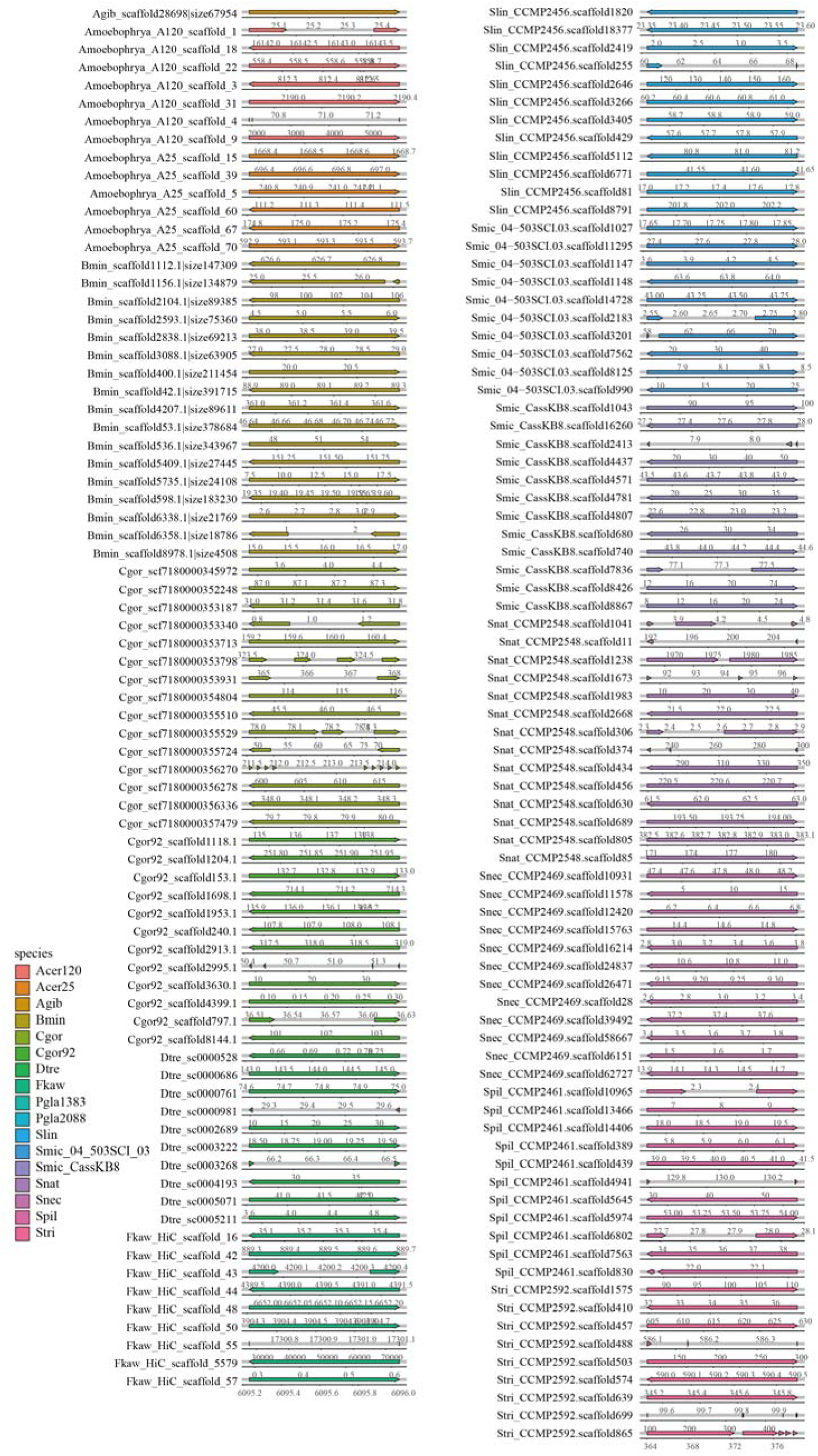
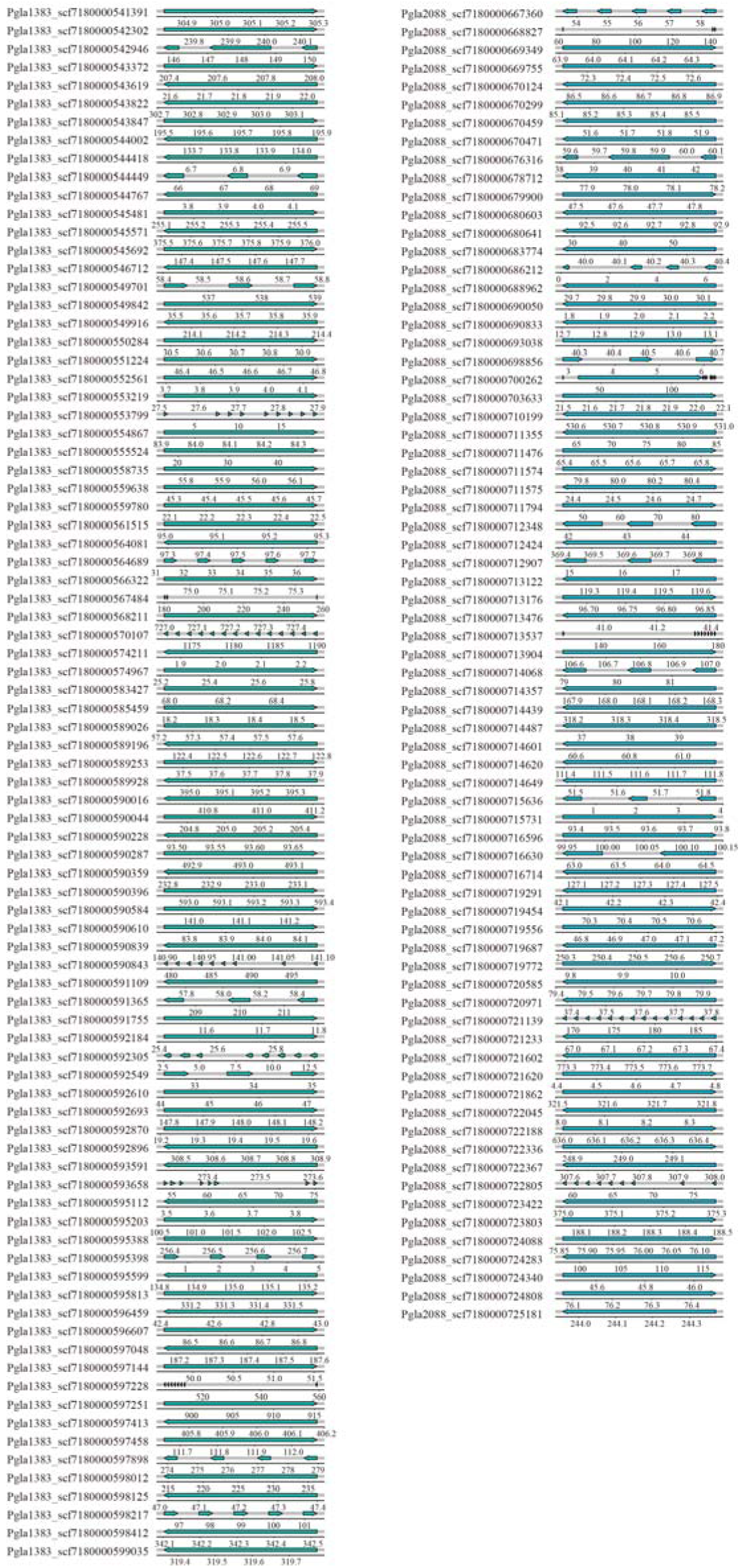
All DVNPs location of dinoflagellates genomes. Arrow to the right indicates that the gene is on the forward strand. The species abbreviation is consistent with the main text.

**Fig S3.**
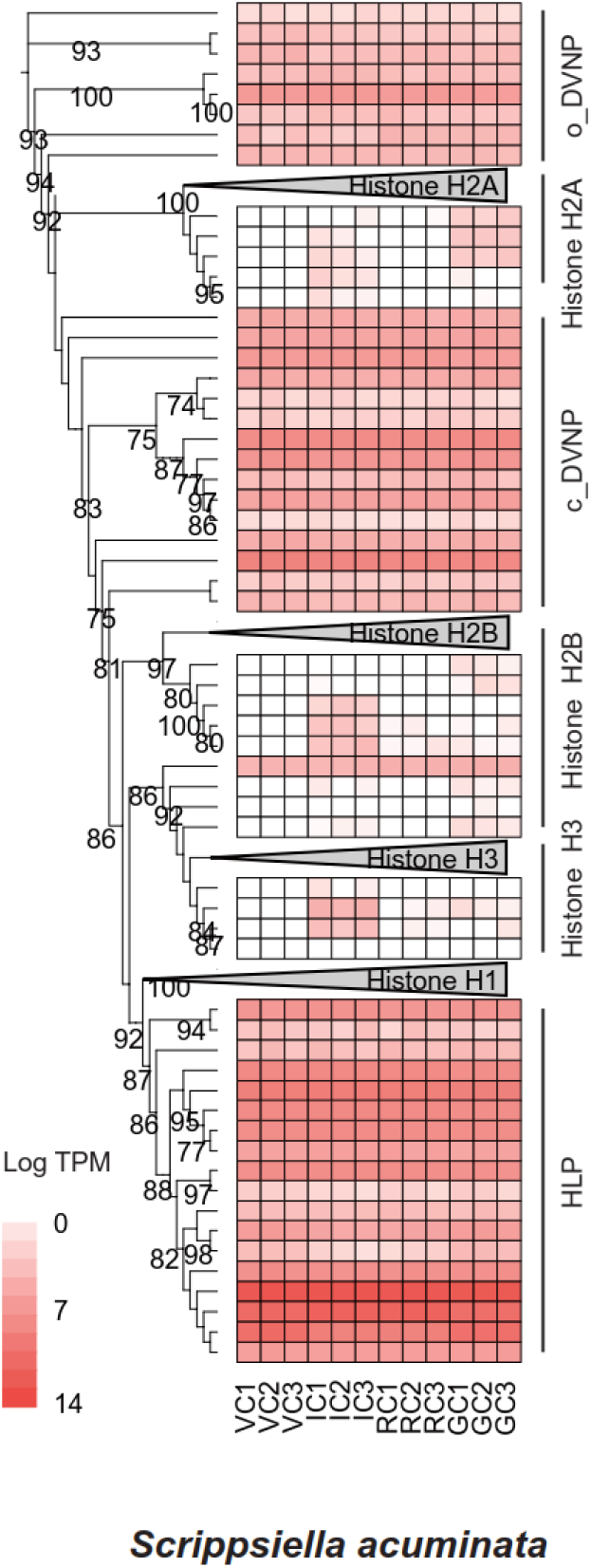
Published data from GenBank’s Sequence Read Archive (SRA) database. PRJNA744760 for *S. acuminate*. Samples of *S. acuminate* representing different cell stages. Vegetative cells (VC), immature cysts (IC), resting cysts (RC) and germinating cysts (GC), ultrafast bootstraps >70% are shown.

**Fig S4.**
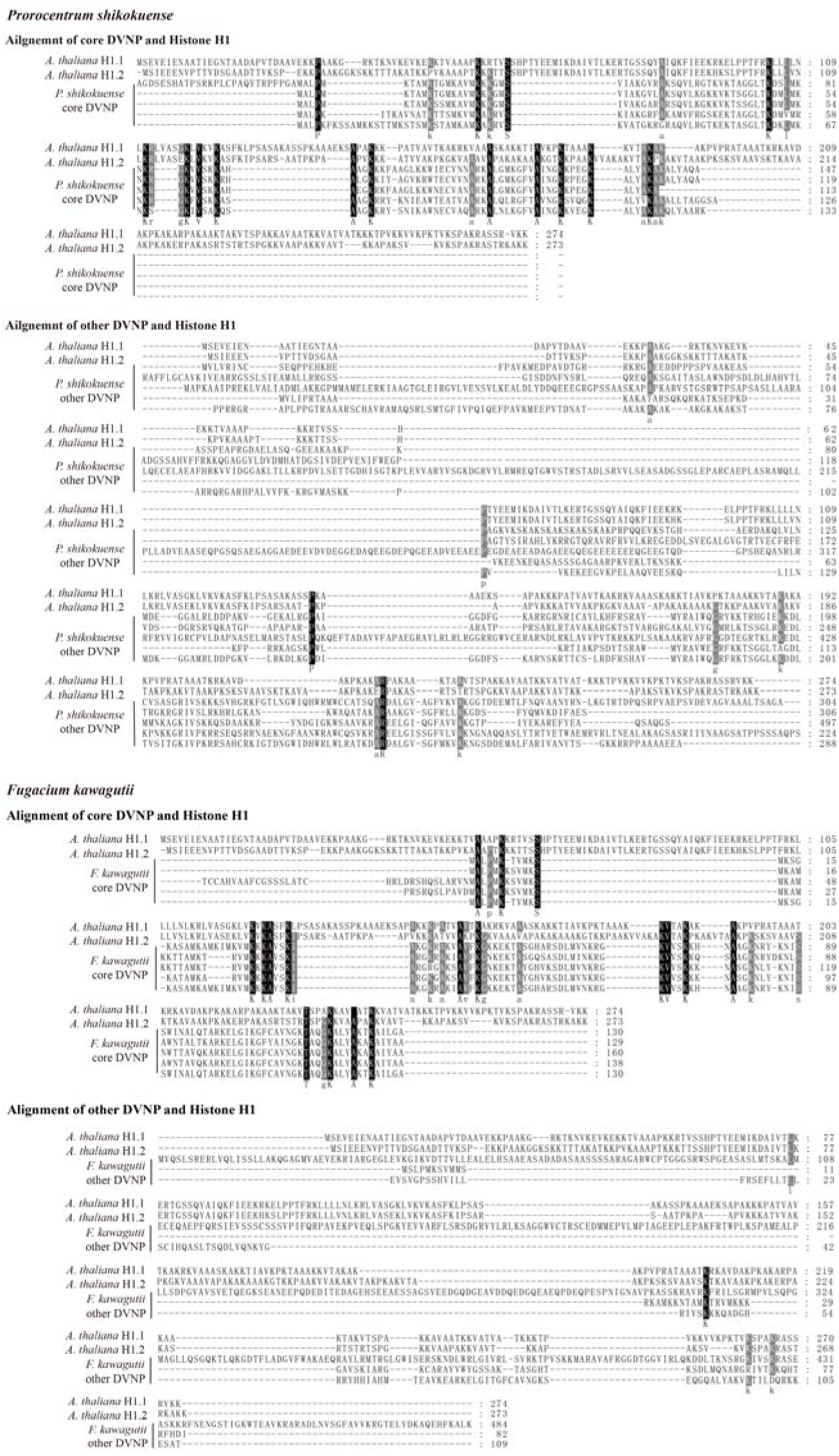
Multiple alignments of *A. thaliana* histone H1 and core DVNP genes of *P. shikokuense* and *F. kawagutii*. Subscript letters indicate conserved residue (conserved percent > 80%). Around 10% identity between core DVNPs and *A. thaliana* histone H1 (9.2% in *P. shikokuense*, 10.3% in *F. kawagutii*) while no more than 2.9% identity between other DVNPs and histone H1.

**Fig S5.**
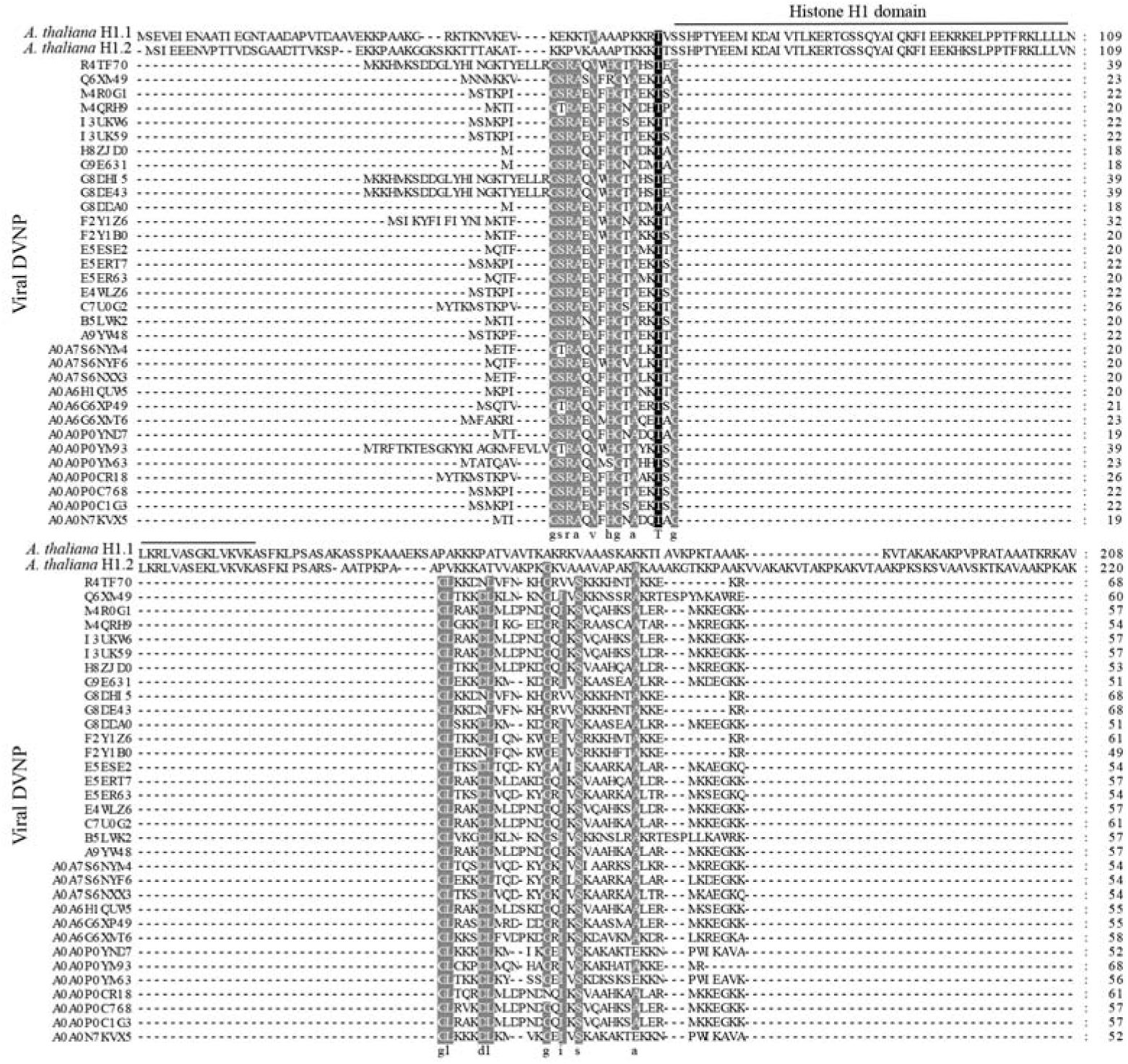
Multiple alignments of *A. thaliana* histone H1 and viral DVNPs. Subscript letters indicate conserved residue (conserved percent > 80%).

**Fig S6.**
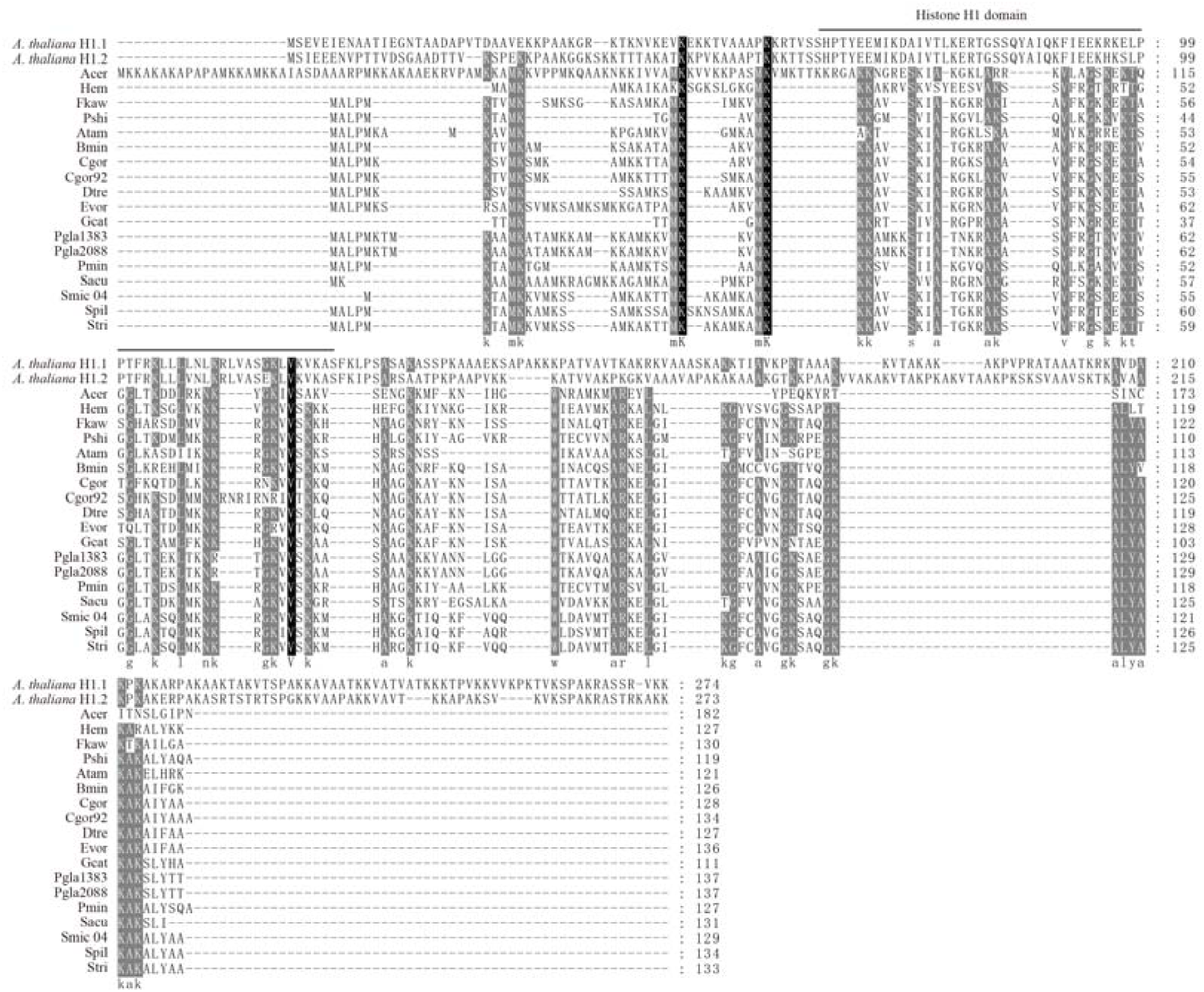
Multiple alignments of *A. thaliana* histone H1 with DVNPs of multi-species in dinoflagellates. Subscript letters indicate conserved residue (conserved percent > 80%).

**Fig S7.**
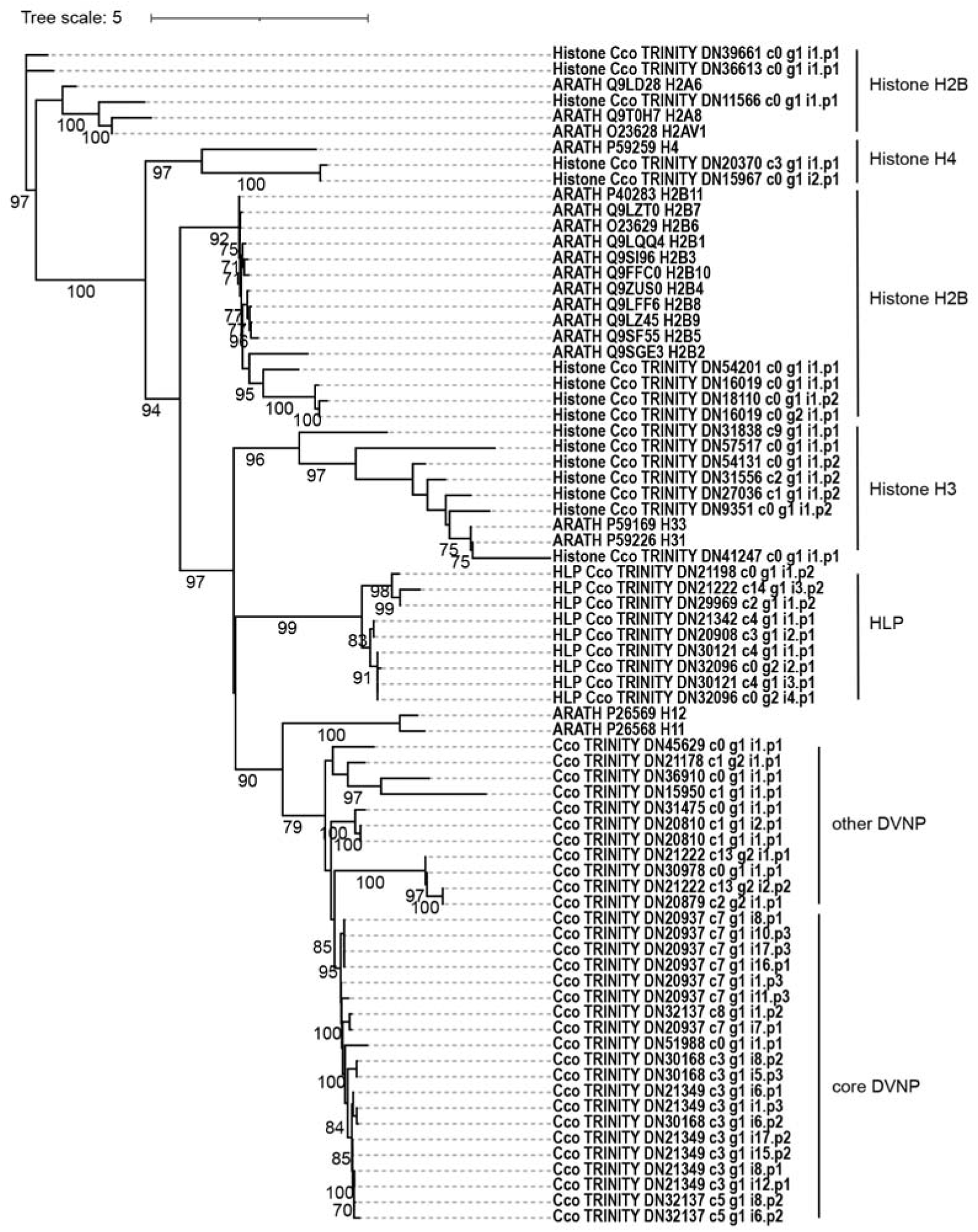
Phylogeny of DVNP, histones and histone-like protein (HLP) of *Crypthecodinium cohniie* (Cco). Ultrafast bootstraps >70% are shown.

**Table S1.** Data sources of published studies.

| Data type | Species | Source |
| --- | --- | --- |
| Transcriptome | *Amphidinium carterae* | MMETSP0398 |
|  |  | MMETSP0399 |
|  |  | MMETSP0258 |
|  |  | MMETSP0259 |
|  | *Prorocentrum minimum*<br>CCMP2233 | MMETSP0267 |
|  |  | MMETSP0268 |
|  |  | MMETSP0269 |
|  |  | MMETSP0053 |
|  |  | MMETSP0055 |
|  |  | MMETSP0056 |
|  |  | MMETSP0057 |
|  | *Noctiluca scintillans* | MMETSP0253 |
|  | *Gymnodinium catenatum* GC744 | MMETSP0784 |
|  | *Alexandrium tamarense*<br>CCMP1771 | MMETSP0382 |
|  |  | MMETSP0384 |
|  |  | MMETSP0378 |
|  |  | MMETSP0380 |
|  | *Alexandrium catenella* OF101 | MMETSP0790 |
|  | *Hematodinium* sp | GenBank: GEMP00000000.1 |
|  | *Karenia mikimotoi* | Luo et al., 2017 |
|  | *Scrippsiella acuminata* | Xiaomei et al., 2022 |
|  | *Prorocentrum shikokuense* | **This study** |
|  | *Cryptocodinium cohnii* | MMETSP0323 |
|  |  | MMETSP0324 |
|  |  | MMETSP0325 |
|  |  | MMETSP0326 |
| Genome | *Perkinsus marinus* | GCA_000006405.1 TIGR |
|  | *Breviolum minutum* | Shoguchi et al., 2013 |
|  | *S. microadriaticum* CassKB8 | González-Pech et al., 2021 |
|  | *S. microadriaticum* 04-503SCI.03 |  |
|  | *S. tridacnidorum* |  |
|  | *S. linucheae* |  |
|  | *S. necroappetens* |  |
|  | *S. natans* |  |
|  | *S. pilosum* |  |
|  | *Cladocopium goreau* | Chen et al., 2023 |
|  | *Cladocopium* C92 | Shoguchi et al., 2018 |
|  | *Fugacium kawagutii* | Li et al., 2020 |
|  | *Effrenium voratum* | PRJEB61191 |
|  | *Durusdinium trenchii* | Shoguchi et al., 2021 |
|  | *Polarella gracialis* CCMP2088 | Stephens et al., 2020 |
|  | *Polarella gracialis* CCMP1383 |  |
|  | *Amoebophrya* A120 | Farhat et al., 2021 |

**Table S2.** The topology tests of *A. thaliana* histone H1 in DVNP tree .

| Tree | bp-RELL<sup>b</sup> |  | P-KH<sup>c</sup> |  | p-SH<sup>d</sup> |  | c-ELW<sup>e</sup> |  | p-AU<sup>f</sup> |  |
| --- | --- | --- | --- | --- | --- | --- | --- | --- | --- | --- |
| H1 in core DVNP | 0.437 | + | 0.511 | + | 1 | + | 0.444 | + | 0.722 | + |
| H1 in other DVNP<sup>a</sup> | 0.0021 | - | 0.0129 | - | 0.101 | + | 0.00251 | - | 0.00304 | - |
| H1 out of DVNP<sup>a</sup> | 0.0062 | - | 0.0256 | - | 0.212 | + | 0.00797 | - | 0.0299 | - |

Plus signs denote the 95% confidence sets. Minus signs denote significant exclusion. All tests performed 10000 resamplings using the RELL method.
<sup>a</sup> hypotheses tree.
<sup>b</sup> bootstrap proportion using RELL method (Kishino et al. 1990).
<sup>c</sup> p-value of one sided Kishino-Hasegawa test (1989).
<sup>d</sup> p-value of Shimodaira-Hasegawa test (2000).
<sup>e</sup> Expected Likelihood Weight (Strimmer & Rambaut 2002).
<sup>f</sup> p-value of approximately unbiased (AU) test (Shimodaira, 2002).

**Table S3.** *P. shikokuense* cultures under four laboratory conditions.

| Culture conditions |  |  | Number of samples |
| --- | --- | --- | --- |
| Light variation | low light | 48h | 3 |
|  | normal light |  | 3 |
|  | high light |  | 3 |
|  | red light | 24h, 96h | 3 |
|  | blue light |  | 3 |
|  | green light |  | 3 |
|  | continuous lighting | 48h | 3 |
|  | dark treatment |  | 3 |
| Nutrient changes | nitrogen limitation |  | 3 |
|  | nitrogen replete |  | 3 |
|  | phosphorus replete |  | 3 |
|  | phosphorus depleted |  | 3 |
| Growth stage | diel | per 2 hours | 3 |
|  | 21:30 |  | 3 |
|  | 0:30 |  | 3 |
|  | 4:40 |  | 3 |
|  | 7:40 |  | 3 |
|  | exponential stage | 5 days after | 3 |
|  | platform stage | 2 weeks after | 3 |
|  | extinction stage | 3 months after | 3 |
| Temperature change | low temperature | 13 °C | 3 |
|  |  | 16 °C | 3 |
|  | normal temperature | 20 °C | 3 |
|  | high temperature | 25 °C | 3 |
|  |  | 28 °C | 3 |

